# 16p11.2 duplication shows early male-biased impacts on reward learning, but NMDA receptor antagonism reduces optimal choice selection in both wildtypes and 16p11.2 duplication

**DOI:** 10.1101/2025.06.28.662127

**Authors:** Dana Mueller, Evan Knep, Angelica Velosa, Erin Giglio, Cathy S. Chen, Sarah R. Heilbronner, R. Becket Ebitz, Patrick E. Rothwell, Nicola M. Grissom

**Author notes:** **to whom correspondence should be addressed**: Nicola Grissom Department of Psychology University of Minnesota, Minneapolis, MN 75 East River Road Minneapolis, MN 55454.

## Abstract

**Rationale:** 16p11.2 duplication is associated with numerous neuropsychiatric conditions at a genome-wide level, including psychosis. Mice modeling 16p11.2 duplication may provide important insights into cognitive risk factors, in particular in reward-guided decision making. NMDAR function has also been implicated in psychosis phenotypes, but whether these phenotypes differ by genetic risk factor is unknown.

**Objectives:** We aimed to: 1) identify sex and genotype differences in early operant training and two-arm spatial restless bandit task performance; 2) examine the effects of an NMDAR antagonist on task performance and strategy across genotypes.

**Methods:** 16p11.2 duplication and wildtype mice completed a series of training schedules of escalating difficulty followed by bandit tasks. MK-801 and saline were administered in alternating sessions prior to later bandit task performance.

**Results:** Large sex differences in early operant training revealed some male-biased impacts of 16p11.2 duplication, contingent on training schedule difficulty. Once on the two-arm spatial restless bandit task, 16p11.2 duplication was no longer a strong contributor to decision making. However, MK-801 decreased the tendency to stay with a rewarded choice, lowered the probability of selecting the highest rewarded option, and decreased the influence of prior outcomes on choice.

**Conclusions:** The male-biased vulnerability in early operant training suggests that strategies for learning early schemas or in novel environments may be impacted by 16p11.2 duplication in males. In contrast, NMDAR are influential in the ability to flexibly switch between choices, and disrupting this function significantly impairs decision making in all animals.

## Introduction

Schizophrenia and psychosis are characterized by not only positive symptoms (such as hallucinations), but also prominent cognitive symptoms for which we lack treatments. Cognitive impairments can impede quality of life, and so understanding the mechanisms behind the symptoms as well as finding treatments for these symptoms is critically needed (Patel et al. 2014; Balu 2016). Decision making deficits in schizophrenia and psychosis are centered around inconsistent choice selections (Geana et al. 2022; Pratt et al. 2021), increased “switching,” or exploration, between choices, and decreased selection of the highest value choice (Gold et al. 2008; Juckel et al. 2006; James A. Waltz et al. 2020; Schlagenhauf et al. 2014; Reed et al. 2020). Understanding such decision making computations can reveal mechanisms of psychosis vulnerability in clinical populations.

Among the gene variants most frequently associated with neuropsychiatric disorders, including schizophrenia and psychosis, is 16p11.2 copy number variation (Yang et al. 2015; Bertero et al. 2018; Wang et al. 2018; Blaker-Lee et al. 2012; Niarchou et al. 2019; Marshall et al. 2016; Rein and Yan 2020). The 16p11.2 region is highly syntenic across species, and cognitive deficits associated with 16p11.2 hemideletion, including male biased operant deficits (N. M. Grissom et al. 2017), have been extensively explored (Yang et al. 2015; Bertero et al. 2018; Wang et al. 2018). In contrast, although 16p11.2 duplication (16p11 Dup) is strongly linked at the level of genome wide significance to schizophrenia and psychosis (McCarthy et al. 2009; Rees et al. 2014; Rein and Yan 2020; Steinberg et al. 2014), the cognitive phenotypes of 16p11 Dup are underexplored. Recent studies provide evidence for social deficits and grooming phenotypes (Rein and Yan 2020) and impaired performance in working memory tasks (Rein and Yan 2020; Rein et al. 2021; Bristow et al. 2020). However, the impacts of 16p11 Dup has not yet been explored in decision making contexts that are strongly linked with human phenotypes.

Another mechanism implicated in cognitive deficits in schizophrenia is NMDAR function, which is known to be altered in patients with schizophrenia (MacDonald and Chafee 2006; Kummerfeld et al. 2020; Zick et al. 2022; Adell 2020; de la Salle et al. 2019; Zick et al. 2018). Specifically, systemic NMDAR blockades prevent coordinated neuronal firing and produce cognitive deficits similar to those seen in schizophrenia (Zick et al. 2022; Vinogradov et al. 2023; Alvarez et al. 2020; Gao et al. 2022; Snyder and Gao 2020). An NMDAR antagonist, MK-801, disrupts rule learning in rat set-shift behavior (Stefani and Moghaddam 2005). NMDAR dysfunction impairs cognition in wildtype mice (Schmack et al. 2021), but it is unknown how pharmacological manipulation of NMDA function interacts with genetic variants associated with risk of psychosis, which can be addressed using mouse models. Testing decision making in a strongly associated genetic variant, 16p11 Dup, offers an opportunity to assess whether there are additive effects of genetic vulnerability and NMDAR dysfunction.

In the current study, we examine reward learning and decision making in male and female 16p11 Dup mice, with and without the NMDAR antagonist MK-801. Building on previous work using bandit and reversal tasks to investigate the flexibility needed in decision making to successfully switch between rules and choices in patients diagnosed with psychosis (Murray et al. 2008; Suetani et al. 2022; Schlagenhauf et al. 2014; Reddy et al. 2016; J. A. Waltz and Gold 2007; McKirdy et al. 2009), the current study employs our two-arm spatial restless bandit task.

We find that early operant training data is consistent with male-biased impacts of 16p11.2 hemideletion, as previously shown (N. M. Grissom et al. 2017). Of these two models for risk factors for psychosis, MK-801 treatment yielded the largest effect, producing a deficit in the ability to flexibly switch between choices to obtain the optimal amount of reward. The cognitive phenotypes demonstrated here may be relevant to some aspects of cognitive impairments in humans with 16p11.2 duplications, helping to provide insight into human neuropsychiatric disorders.

## Methods

### Subjects

Male 16p11.2 duplication (16p11 Dup) mice (stock # 013129) and female wildtype mice (stock # 101043) were obtained from Jackson Laboratories and bred in order to generate mice of both genotypes for the experiments. Mice were housed in groups of 2-5 of mixed genotypes. Data included in this manuscript are combined from two cohorts that were run consecutively (total n: 16 male 16p11 Dup, 23 male wildtype, 21 female 16p11 Dup, and 26 female wildtype mice). The first cohort (8 male 16p11 Dup, 13 male wildtype, 9 female 16p11 Dup, and 11 female wildtype) completed 20 sessions of the two arm restless bandit task. The second cohort (8 male 16p11 Dup, 10 male wildtype, 12 female 16p11 Dup, and 15 female wildtype) completed 20 sessions of the two arm restless bandit task and 12 sessions of interleaved saline and MK-801 manipulation. *Transnetyx* was used to confirm genotypes both before and after behavioral experiments. Experimenters were blind to genotype while handling the mice. Mice began behavioral training at approximately 12 weeks of age. Mice had free access to water and were food restricted to no lower than 85% of their free-feeding body weight. Behavioral testing took place Monday to Friday, and on Friday, mice had free access to home chow. Colony rooms were temperature controlled (20.5°C; 69°F) and on a light-dark cycle of 12 hours (08:00-16:00), where animals were tested during the dark period. All animals were cared for according to the guidelines of the National Institutes of Health and the University of Minnesota IACUC approval.

### Apparatus

Preliminary training and behavioral testing were carried out in the same Bussey-Saksida touchscreen chambers for all mice throughout the present study (Lafayette Instrument Company, Lafayette, IN) which are sensitive to continuous and rapidly repeated touches in the same location and across the entirety of the screen (Heath, Bussey, and Saksida 2015). The sound-reducing chamber includes black plexiglass walls, a stainless steel grid floor, and a removable lid. Each chamber contained an automated food dispenser where 50% water-diluted vanilla Ensure was delivered. An opaque mask covered the screen with two response apertures for the two choices in the two-arm spatial restless bandit task. These two response apertures display the trial stimulus (two illuminated squares - left and right choices) and are within a mouse’s reach from the floor of the chamber.

### Behavioral Training Program

*Chamber acclimation.* Prior to the first day in the operant chamber (“Day 0”), training mice were pre-exposed to vanilla Ensure in a bottle overnight allowing them familiarity with the reward prior to receiving the reward for the first time during a training schedule. *Day 0* is a single habituation day in the operant chamber where free reward (50µL; vanilla Ensure) is dispensed at the very beginning of a 30 minute exposure to the operant chamber.

*Initial Touch Training.* This training trains mice to use the touchscreen and reinforces animals with a larger reward for interacting with the touchscreen. A free reward (7µL) is given every 30 seconds; however, if a mouse touches any coordinates within the choice aperture containing a random image it will get an additional immediate reward dispensed which is 3x the amount of the free reward (21µL; **Figure 1a**). This training schedule lasts 30 minutes. Mice remain on this schedule until they complete a minimum of 30 trials in 30 minutes for two consecutive days.

**Figure 1.**
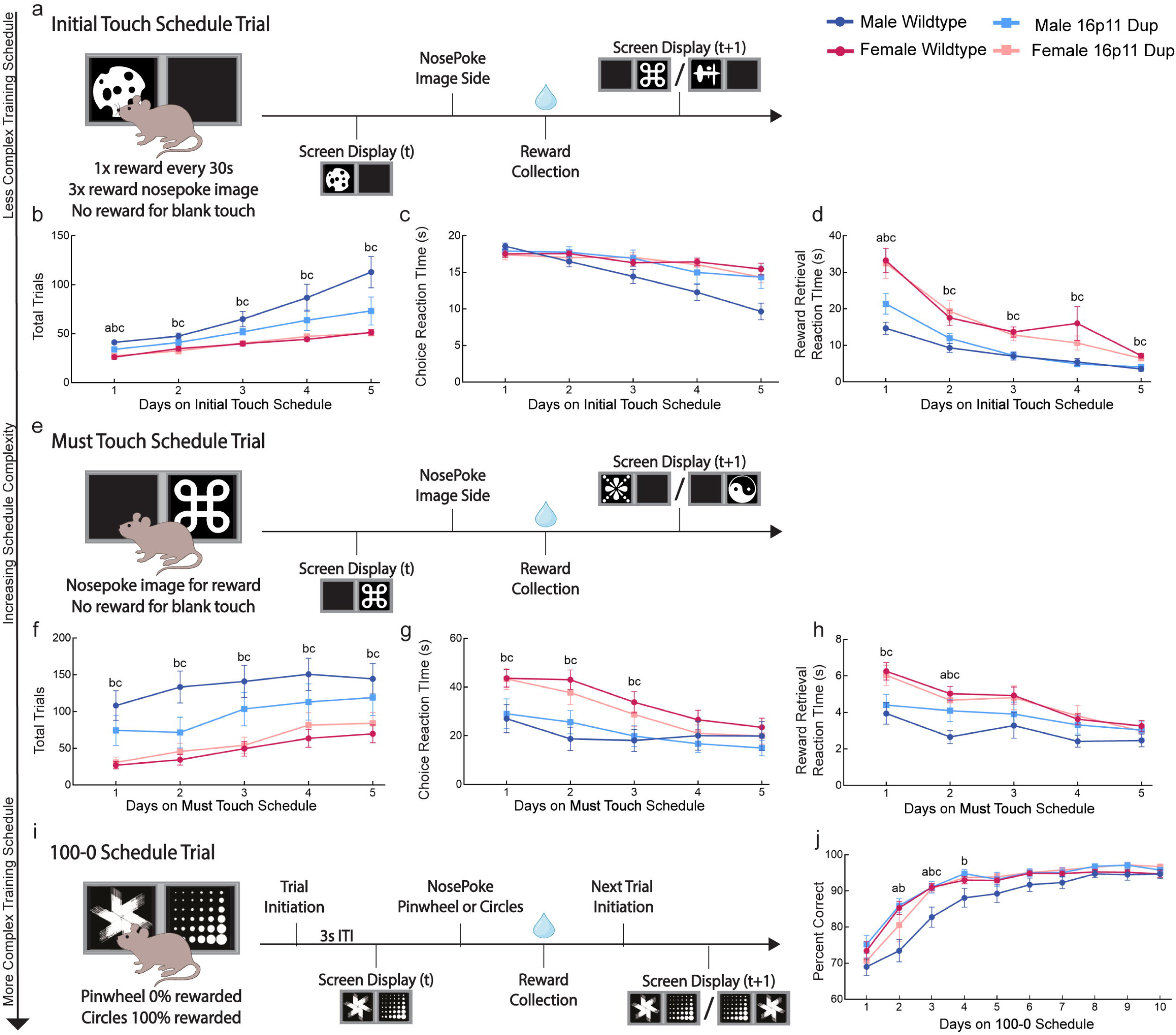
16p11.2 duplication prevents male-typical engagement with operant training, with no impact on females. a) Schematic illustrating possible screen display and trial events for *Initial Touch* training schedule. b) Total trials displayed across the first 5 days of *Initial Touch* training split by sex and genotype. Male wildtype mice had significantly more total trials. Planned pairwise comparisons revealed that male wildtypes completed more trials than male 16p11 dup on day 1 (p = 0.045; indicated by “a”) and all females on all days (all p ≤ 0.006; female wildtype indicated by “b”; female 16p11 Dup indicated by “c”). c) Choice reaction time displayed across the first 5 days of *Initial Touch* training split by sex and genotype. Male wildtype mice had significantly reduced choice reaction times. d) Reward retrieval reaction time displayed across the first 5 days of *Initial Touch* training split by sex and genotype. Males had significantly reduced reward retrieval reaction times. Planned pairwise comparisons revealed that male wildtypes collected reward faster than male 16p11 dup on day 1 (p = 0.031) and all females on all days (all p ≤ 0.003). e) Schematic illustrating possible screen display and trial events for *Must Touch* training schedule. f) Total trials displayed across the first 5 days of *MustTouch* training split by sex and genotype. Male wildtype mice had significantly more total trials. Planned pairwise comparisons revealed that male wildtypes completed more trials than all females on all days (all p ≤ 0.050). g) Choice reaction time displayed across the first 5 days of *MustTouch* training split by sex and genotype. Males had significantly reduced choice reaction times. Planned pairwise comparisons revealed that male wildtypes chose faster than all females on days 1, 2, and 3 (all p ≤ 0.014). h) Reward retrieval reaction time displayed across the first 5 days of *MustTouch* training split by sex and genotype. Males wildtype mice had significantly reduced reward retrieval reaction times. Planned pairwise comparisons revealed that male wildtypes collected reward faster than male 16p11 dup on day 2 (p = 0.049) and all females on days 1 and 2 (all p ≤ 0.010). i) Schematic illustrating possible screen display and trial events for *100-0 deterministic* training schedule. j) Percent Correct displayed across the first 10 days of *100-0 deterministic* training split by sex and genotype. Male wildtype mice had significantly less percent correct. Planned pairwise comparisons revealed that male wildtypes have a lower percent correct than male 16p11 dup on day 2 and 3 (all p < 0.009), than female wildtype on days 2, 3, and 4 (all p ≤ 0.037), and than female 16p11 dup on day 2 (p = 0.021). For simplicity of visualization, all plots are averages across trials and mice within a particular session. Red circles indicate female wildtype, blue circles indicate male wildtype, light blue squares indicate male 16p11 Dup, and orange squares indicate female 16p11 Dup mice. Significance throughout this figure is represented in the following way: “a” indicates male wildtype significantly different from male 16p11 Dup, “b” indicates male wildtype significantly different from female wildtype, “c” indicates male wildtype significantly different from female 16p11 Dup. Error bars represent standard error of the mean (SEM).

*Must Touch Training.* This training requires mice to use the touchscreen to gain rewards. There is no longer a free reward every 30 seconds, but rather if a mouse touches any coordinates within the choice aperture containing a random image, immediate reward is dispensed (7µL; **Figure 1e**). This training schedule lasts 30 minutes. Mice remain on this schedule until they complete a minimum of 30 trials in 30 minutes for two consecutive days.

*Pairwise Must Initiate Training.* This training trains mice to initiate a trial by entering the reward port. At this stage of training mice have learned to touch the screen after it has been lit to gain a reward (7µL). The reward is followed by a three second inter-trial-interval (ITI) and then a light cue at the reward point is turned on to signal the mouse to enter the reward port to initiate a new trial (**Supplemental Figure 1b**). This training schedule lasts 30 minutes. Mice remain on this schedule until they complete a minimum of 30 trials in 30 minutes for two consecutive days.

*Pairwise Punish Incorrect Training.* This training trains the precision of touchscreen response to the lit-up image only. If a mouse wrongly touches the unlit choice aperture (e.g. left side instead of right side), the operant chamber house light will blink for two seconds and there will be a ten second timeout as punishment. If the mouse correctly touches the lit portion of the screen, they are rewarded (7µL), followed by a three second inter-trial-interval (ITI) and the mouse must enter the reward port to initiate a new trial (**Supplemental Figure 1g**). This training schedule lasts 60 minutes or until 200 trials. Mice remain on this schedule until they complete 200 trials for two consecutive days.

*100-0 Deterministic Learning Training.* This deterministic training schedule is the first value-based decision making training that requires a mouse to choose between two images and learn about the correct image from feedback. One image (circles) is rewarded 100% of the time and the other image (pinwheel) is rewarded 0% of the time with no punishment timeout (**Figure 1i**). The rewarded image switches between left and right side, but is always rewarded regardless of spatial location. This training schedule lasts 120 minutes or until 250 trials. Mice remain on this schedule until they consistently complete 250 trials with a percent correct greater than 85 percent.

*Probabilistic Learning Training.* This training consists of a series of probabilistic reversal learning schedules. 90-10 spatial training requires a mouse to choose between left and right side (identical visual cue), where one side is rewarded (7µL) 90% of the time, and the other side is rewarded 10% of the time. 80-20 and 70-30 spatial trainings are the same with 80% vs. 20% rewarded options and 70% vs. 30% rewarded options. The reward probability associated with the left and right side reverse based on choice matching probability of reward, e.g. 90-10 reversal occurs after the high-value choice is chosen 9 out of the last 10 trials. Mice experience one day o*f 90-10 Spatial*, followed by one day of *80-20 Spatial*, followed by two days of *70-30 Spatial*, just prior to transition to bandit schedules. The purpose of this training is to adapt mice to a stochastic and changing environment, prior to the restless bandit task. These training schedules last 120 minutes or until 300 trials.

*Restless Bandit Behavioral Paradigm.* After completing the pre-training operant schedules, mice began two-arm spatial restless bandit schedules. During experiment mice complete sessions of the two-arm restless bandit task Tuesday-Friday and *100-0 Deterministic Learning Training* is completed every Monday. Mice did not run on weekends. In our bandit task, mice must decide between two spatial choices on a touchscreen (**Figure 2a**). On every trial there is a 10% chance of the reward probability associated with each arm (choice) increasing or decreasing by 10%. The reward contingency is always stochastic, which means the reward probability cannot go down to 0% or up to 100%. Each day of bandit consisted of a new walk of independent and randomly changing probabilities (**Figure 2b**) so that the task could not be remembered from day to day. This schedule lasts 120 minutes or until 300 trials.

**Figure 2:**
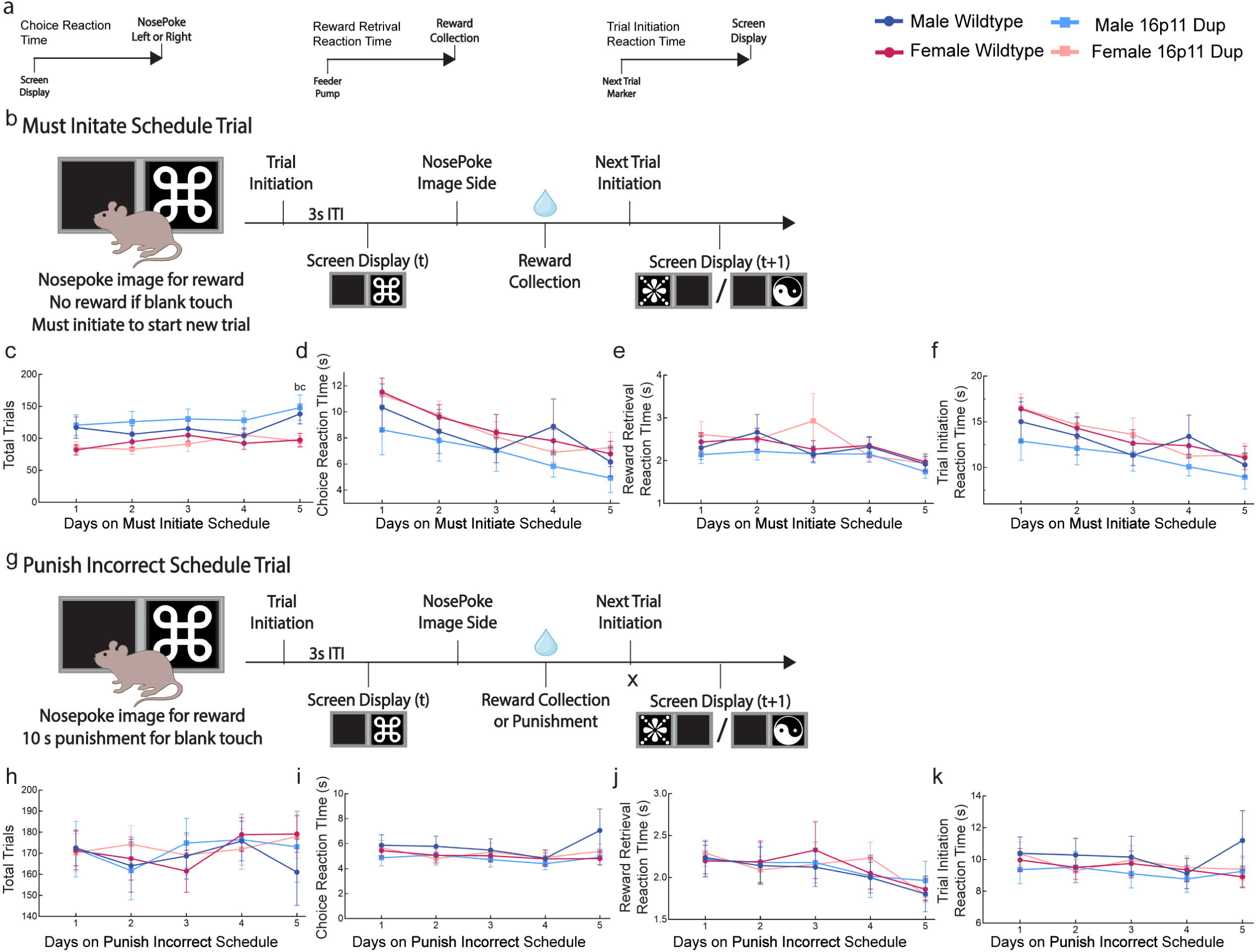
Restless bandit performance reveals 16p11.2 duplication enhances outcome sensitivity. a) Schematic depicting the timeline of a single two-arm spatial restless bandit trial. White squares indicate left/right spatial choice. b) Schematic depicting the hidden Markov model (HMM) and labeling explore trials along an example two-arm spatial restless bandit probability walk. Orange traces indicate the probability and choices of left side touches. Blue traces indicate the probability and choices of right side touches. Gray shaded regions indicate HMM labeled explore trials. c) Total trials displayed across bandit sessions 1-20 split by sex and genotype. Red circles indicate female wildtype, blue circles indicate male wildtype, light blue squares indicate male 16p11 Dup, and orange squares indicate female 16p11 Dup mice. Males completed significantly fewer total trials. d) Percent Reward - Chance split by genotype and sex. No significant differences between groups. Red indicates female wildtype, blue indicates male wildtype, light blue indicates male 16p11 Dup, and orange indicates female 16p11 Dup mice. e) Outcome weight split by genotype and sex. 16p11 Dup mice had significantly increased outcome weight compared to wildtype mice. Red indicates female wildtype, blue indicates male wildtype, light blue indicates male 16p11 Dup, and orange indicates female 16p11 Dup mice. For simplicity of visualization, all plots are averages across trials and sessions, so that each individual data point plotted represents the overall average for a mouse. Significant throughout this figure is represented in the following way: * p value less than 0.05. Error bars represent standard error of the mean (SEM).

### Drug Administration

To assess the effect of NMDAR activity on bandit task strategy and response accuracy, we employed MK-801 (also known as dizocilpine), a non-competitive NMDAR antagonist. MK-801 was fully dissolved in 0.9% saline and protected from light. Animals received an intraperitoneal (i.p) injection of drug or saline (control) at an injection volume of 5ml/kg immediately before entering the chambers, allowing for efficacy over the 2 hours of bandit task testing. Drug and saline were administered in alternating sessions using a within-subjects design, so that every animal completed 12 total sessions (6 saline/6 MK-801 sessions). Interleaving control and drug sessions allowed us to account for potential effects of repeated drug administration and long-term changes.

Dosage chosen was based on previous studies on the effect of MK-801 on cognitive functions (Kruk-Slomka et al. 2016; Bygrave et al. 2016; Mandillo et al. 2003; Uehara et al. 2010; Stefani and Moghaddam 2005; Li et al. 2015). Preliminary studies showed that higher doses (e.g., 1 mg/kg) produced strong psychomotor activation that could interfere with task performance (data not shown). A 0.1 mg/kg dose was chosen with the intention of implementing the lowest dose necessary to produce alterations in cognitive functions including decision-making, learning, and exploration while minimizing effects on locomotion (Bradford et al. 2010; Kruk-Slomka et al. 2016; He et al. 2020; Kraeuter et al. 2020; Gray et al. 2009; Bygrave et al. 2016). The effect of MK-801 was only tested on the second cohort (8 male 16p11 Dup, 10 male wildtype, 12 female 16p11 Dup, and 15 female wildtype).

### Data Analysis

All raw touchscreen choices were collected by Lafayette ABET software allowing for quantification of total trial counts, rewards earned, response times, trial initiation times, etc. per mouse and session.

*Choice Reaction Time.* To determine latency to respond across training and two-arm spatial restless bandit task schedules we calculated choice reaction time in seconds. Choice response time was calculated as the time elapsed between screen display onset and the time when the nosepoke to the left or right choice aperture was completed (**Supplemental Figure 1a**).

*Reward Retrieval Reaction Time.* To determine latency to retrieve reward across training and two-arm spatial restless bandit task schedules we calculated reward retrieval reaction time in seconds. Reward retrieval reaction time was calculated as the time elapsed between reward delivery (feeder pump) and reward collection (**Supplemental Figure 1a**).

*Trial Initiation Reaction Time.* To determine latency to initiate a new trial across training and two-arm spatial restless bandit task schedules we calculated trial initiation reaction time in seconds. Trial initiation reaction time was calculated as the time elapsed between completed trial (next trial increment) and the next trial screen display (prompted by magazine entry; **Supplemental Figure 1a**). Trial initiation reaction time is not a measure that can be calculated for *Initial Touch Training* and *Must Touch Training* since the mice do not learn to initiate until *Pairwise Must Initiate Training*.

*P(Best) Analysis.* The probability that a mouse chooses the choice with the objectively highest reward probability on a given trial. This provides a measure of choice optimality. When the two arms (choices) match in probability, then either choice is considered “best” for calculation purposes.

*Outcome Weight Analysis.* We quantified the relative impact of negative versus positive outcomes on the probability of switching choices within a session for each mouse. Outcome weight is calculated as the probability of shifting choices given no reward minus the probability of shifting choices given reward, all divided by the overall probability of shifting.

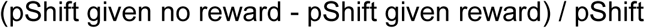

*Touchscreen Analysis.* Each touchscreen aperture represents the x,y coordinates of each response an animal makes on the screen from IR beam technology. The IR emitters are positioned along two sides of the screen and IR receivers are positioned along the other two sides of the screen. With the touchscreen data we calculated Distance from the center of the screen, Euclidean distance, and Mahalanobis distance (Mueller et al. 2025). *Euclidean distance* (Walther et al. 2016; Ebitz and Hayden 2021) is the hypotenuse between two points with (x,y) coordinates that were successive, from the same choice aperture (left/right), and within the same HMM decision state (explore/exploit). *Mahalanobis distance (Walther et al. 2016; Ebitz and Hayden 2021)* is the distance of each touch coordinate from the centroid (central point) of the data cluster.

*Hidden Markov Model.* Because the spatial bandit task is independently and randomly changing, mice are required to evaluate the benefits of exploration and exploitation as they respond to cues to maximize reward. A hidden Markov model (HMM) was used to determine when animals were exploring or exploiting their options in the restless bandit task **(Figure 2b)**. P(exploration) is the probability of mouse exploration between choices throughout the task. The HMM identifies whether mice have entered a state of exploration or a state of exploitation. Chen et al. (2021) has shown performance differences between states such as differences in reaction time and influences of sex on learning during exploration periods. The state of the mouse may change throughout the session based on the comparison of multiple successive trials. For example a mouse may start off in an exploratory state until a rewarding option is identified and then enter an exploitative state for a number of trials. Each of the 300 total trials can therefore be grouped and labeled as exploratory or exploitative, and analyses were performed to compare distance and time between successive touches across states, as well as measures of spatial decision making. **Figure 2b (left)** shows that a jump can not be made from exploiting the left choice to exploiting the right choice, but that each mouse must enter a period of exploration as part of the transition to exploiting an alternative choice. **Figure 2b (right)** then illustrates how trials can be labeled as explore across the restless bandit 300 trial probability walk (shaded in gray). For validation of this model please see (Chen, Knep, et al. 2021).

*Mutual Information Analysis.* We quantified the extent to which choice history was informative about current choices as the conditional mutual information between the current choice and the previous choice based on the reward outcome of the previous trial where the set of choice options represented the two options (left/right) (Chen, Ebitz, et al. 2021; Chen, Knep, et al. 2021; Leao, Fragoso, and Ruffino 2017; Wyner 1978). The current choice is represented by (C_t_). The last choice is represented by (C_t-1_). Reward outcome of the previous trial is represented by (R).

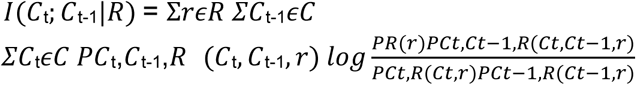

*Generalized linear mixed models.* Data was analyzed with custom Python and GraphPad Prism 10 scripts. Generalized linear mixed models (Python package pymer4) were used to determine state, sex, drug, and genotype differences, unless otherwise specified (Jolly 2018). P values were compared against a standard α = 0.05 threshold. Significant throughout this paper is represented in the following way: * p value less than 0.05. For the portion of the experiment without drug manipulation, sample size is n = 16 male 16p11 Dup, 23 male wildtype, 21 female 16p11 Dup, and 26 female wildtype mice. For the MK-801 portion of the experiment, sample size is n = 8 male 16p11 Dup, 10 male wildtype, 12 female 16p11 Dup, and 15 female wildtype. No animal was excluded from the experiment. All statistical tests used and statistical details were reported in the results. For simplicity of visualization, all bar plots are averages across trials and sessions, so that each individual data point plotted represents the overall average for a single mouse.

Winning models were selected using a stepwise GLMM approach starting by including sex, genotype, state (for touchscreen analysis only), and drug (for MK-801 analysis only) as categorical fixed variables and individual mouse identity and cohort (where relevant) as categorical random variable - as well as all pairwise interactions between all included variables. During the model selection process, each child model was created by dropping one variable or interaction from the parent model and choosing the model with the lowest AIC until no drops in AIC were observed without completely dropping significant main effects. Gamma distribution was used for all response time statistics, binomial distribution was used for data with a ceiling effect such as total trials, and normal distribution was applied for all other statistical measures. All models with multiple main effects following the model selection process included interaction terms, however this term is not reported unless statistically significant.

*Analysis of Variance*. For post-hoc analyses we conducted ANOVAs within each day to determine within-day group differences. ANOVA models were run using the Python Scipy package with genotype and sex as independent variables.

*Mann Whitney U.* Because our data was not normally distributed, for post-hoc analyses we conducted Mann Whitney U across male wildtype, female wildtype, male 16p11 Dup, and female 16p11 Dup groups on individual days to assess whether the distributions of any two groups significantly differed. Mann Whitney U models were run using the Python Scipy package with genotype and sex as independent variables.

## Results

16p11.2 duplication (16p11 Dup) is one of the most common copy-number variants significantly associated with psychosis in human patients at the genome-wide level. To understand the behavioral and cognitive phenotypes associated with this duplication we took advantage of serial operant training data and the two-arm spatial restless bandit task recorded in touchscreen operant chambers. Due to the randomly changing probability throughout the two-arm spatial restless bandit task, mice must demonstrate cognitive flexibility by continually learning across 300 trials rather than just at the beginning of the session.

### 16p11.2 duplication prevents male-typical engagement with operant training, with no impact on females

The first training schedule mice experience is *initial touch training,* which provides the first opportunity to assess operant engagement and motivation. In this schedule, mice receive small, free rewards every 30 seconds regardless of engagement, but are reinforced with 3 times the milkshake volume for a nosepoke response on a random image (**Figure 1a**). We tabulated the number of nosepoke responses animals performed over and above free rewards collected. There is a clear trend in mice completing more overall trials than female mice across the first five days on this schedule (**Figure 1b**, GLM p = 0.085; ꞵ_sex_ = 1.740). A planned comparison approach revealed significant group differences in total trials on Day 1 (p < 0.001; f = 10.988), Day 2 (p = 0.001; f = 5.743), Day 3 (p < 0.001; f = 6.915), Day 4 (p = 0.002; f = 5.501), and Day 5 (p < 0.001; f = 8.684). These differences were strongest between male wildtypes and females, but are also apparent between male 16p11 Dup and wildtype males on Day 1.

Along with completing more overall trials in *initial touch*, male mice also had faster reaction times. With this schedule the mice must choose a choice by interacting with the touchscreen in the operant chamber. GLM revealed no significant effects of sex or genotype on choice reaction times (**Figure 1c**). However, male mice collected reward significantly faster than female mice across all five days (**Figure 1d**, p < 0.001; ꞵ_sex_ = 4.492) suggesting a higher motivation to gain reward on this particular task. A planned comparison approach revealed significant group differences in reward collection time on Day 1 (p < 0.001; f = 8.655), Day 2 (p = 0.002; f = 5.380), Day 3 (p < 0.001; f = 7.191), Day 4 (p = 0.025; f = 3.293), and Day 5 (p < 0.001; f = 7.654). These differences were strongest between male wildtypes and females, but are also apparent between male 16p11 Dup and wildtype males on Day 1.

The second training schedule mice experience is *must touch training*, which like *initial touch training*, provides an opportunity to assess operant engagement and motivation, without punishment for off-target responses. In this schedule, mice are reinforced for a nosepoke response on a random image, but no longer receive an automatic reward every 30 seconds (**Figure 1e**). There was a trend for male mice completing more overall trials than female mice across the first five days on this schedule (**Figure 1f**, p = 0.108; ꞵ_sex_ = 1.625). Observation of the graph shows large differences between males and females, but the GLM model fit that excludes the interaction between sex and genotype (and revealed a significant main effect of sex) had a less good model fit (AIC = 4407.2457) than the model we used, which includes the interaction term (AIC = 4397.8856). A planned comparison approach revealed significant group differences on Day 1 (p < 0.001; f = 8.087), Day 2 (p = 0.001; f = 8.804), Day 3 (p < 0.001; f = 7.300), Day 4 (p = 0.005; f = 4.654), and Day 5 (p = 0.011; f = 3.980) between male wildtypes and females of both genotypes. 16p11 Dup males were not significantly different from wildtype males, likely due to variability.

On the *must touch training* schedule, along with completing more overall trials, male mice also had faster reaction times. There was a trend for male mice responding quicker than female mice across the five days on this schedule (**Figure 1g**, p = 0.103; ꞵ_sex_ = 1.630), with planned comparisons revealing this effect was strongest on Day1 (p = 0.022; f = 3.403), Day2 (p = 0.001; f = 6.199), and Day 3 (p = 0.048; f = 2.747). A sex difference in reward collection was also seen but decreased over time, (**Figure 1h**, p < 0.001; ꞵ_sex_ = 3.613) on Day 1 (p = 0.005; f = 4.562) and Day 2 (p = 0.001; f = 5.982).

The third training schedule mice experience is *must initiate training,* which in addition to selecting the random image to receive reward, mice must now initiate a new trial by entering the reward port a second time following reward collection, which will trigger the next screen display (**Supplemental Figure 1b**). Male mice completed more overall trials than female mice across the first five days on this schedule (**Supplemental Figure 1c**, p = 0.006; ꞵ_sex_ = 2.817). A planned comparison approach revealed significant group differences in total trials on Day 5 (p = 0.013; f = 8.684). Male mice also had faster reaction times. Male mice make choices significantly faster than females across the first five days of this schedule (**Supplemental Figure 1d**, p = 0.001; ꞵ_sex_ = 3.365). This is less visible in reward retrieval (**Supplemental Figure 1e**, p = 0.101; ꞵ_sex_ = 1.640) suggesting sex differences in motivation may be shrinking with extended training. With this schedule the mice must now initiate a new trial by entering the reward port opposite the touchscreen in the operant chamber. The time it takes to initiate this new trial can be calculated as trial initiation reaction time (**Supplemental Figure 1a**). Male mice initiate the next trial significantly faster than female mice across all five days (**Supplemental Figure 1f**, p = 0.004; ꞵ_sex_ = 2.916).

The fourth training schedule mice experience is *punish incorrect training* where, building upon all aspects of previous training schedules, mice must now face punishment via a 10 second time-out period if they make a wrong choice. A wrong choice happens when the mouse nosepokes the choice aperture without the random image display (**Supplemental Figure 1g**). Male mice no longer complete more overall trials than female mice across the first five days (**Supplemental Figure 1h**, p = 0.976; ꞵ_sex_ = -0.031) suggesting that the groups converge in performance at this level of training. There are no longer significant sex differences in choice reaction time (**Supplemental Figure 1i**), reward retrieval reaction time (**Supplemental Figure 1j**), or trial initiation reaction time (**Supplemental Figure 1k**). Together this suggests that with extended training on basic operant schedules, behavior across all sex and genotype groups converged.

The last training schedule mice experience is a *100-0 deterministic training* where, for the first time, two images are presented simultaneously - one in each choice aperture. One image is rewarded 100% of the time and the other 0% of the time, regardless of which side of the screen the images are presented on. This training schedule is the most complex, requiring mice to initiate a trial at the reward port, nosepoke the screen display to choose the image with the highest reward probability, collect reward, and then finally enter the reward port again to initiate the next trial **(Figure 1i**). The model used included an interaction term between sex and genotype, which was significant (**Figure 1j**, p = 0.032; ꞵ_sex/genotype_ = -2.179) indicating that the sex effect is driven by the low percent correct achieved by male wildtype mice specifically, compared to all other groups. A planned comparison approach revealed significant group differences in percent correct on Day 2 (p = 0.017; f = 3.581), Day 3 (p = 0.002; f = 5.329), and Day 4 (p = 0.025; f = 3.295), suggesting that male wildtype mice struggle to learn the parameters of the deterministic task early on but level with the other groups in performance around Day 5. GLMMs revealed no significant differences in sex or genotype for total trials, choice reaction time, reward retrieval reaction time, or trial initiation reaction time analyses (figures not shown).

Together these results suggest that while male mice, most strongly male wildtype mice, are highly engaged in early training schedules, but sex differences are mitigated as schedule complexity increases. Most surprisingly, in the *100-0 deterministic* schedule the male wildtype mice struggle to match male 16p11 Dup, female wildtype, and female 16p11 Dup groups in percent correct in early training. While we might have expected the 16p11 Dup genotype to impair learning and cognitive flexibility, instead we see that male 16p11 Dup mice perform as well as both other female groups, although sometimes differently than wildtype males.

### Restless bandit performance reveals 16p11.2 duplication enhances outcome sensitivity

Bandit and reversal tasks have been used to investigate the flexibility needed in decision making to successfully switch between rules and choices in patients diagnosed with psychosis (Murray et al. 2008; Suetani et al. 2022; Schlagenhauf et al. 2014; Reddy et al. 2016; J. A. Waltz and Gold 2007; McKirdy et al. 2009). Here we find that the 16p11 Dup does not impact the ability to flexibly switch between probabilistically rewarded choices, but does increase the influence of outcomes on choices compared to wildtypes.

In our two-arm spatial restless bandit task (**Figure 2a**), the probability of reward of each left and right choice changes independently and randomly of the other, with a 10% chance of probability change on each trial (**Figure 2b**, example probability walk). The unpredictability of this task encourages mice to continually learn and survey their choices, exploring to find the best option and exploiting a good rewarding option across a 300 trial session. Explore and exploit trials were labeled using an HMM approach (Chen, Knep, et al. 2021; Ebitz, Albarran, and Moore 2018) where each trial was defined as either an explore choice or an exploit choice on the left or the right (**Figure 2b**; gray is explore). Mice explore between the two choices or exploit the high-value choice throughout each session in order to maximize reward.

There is no overall sex or genotype difference in *two-arm spatial restless bandit* total trials (**Figure 2c**). There was also no difference in average reward-chance performance (**Figure 2d**). However, we did see a significant effect of 16p11 Dup on outcome weight (**Figure 2e**, p = 0.015; ꞵ_genotype_ = -2.496) which measures the probability of switching choices because of negative outcomes as a proportion of total switching of choices. These data suggest that 16p11 Dup are more sensitive to outcomes in driving switch behavior.

There was no overall sex or genotype difference in restless bandit percent Win-Stay (**Supplemental Figure 2a**) or percent Lose-Shift (**Supplemental Figure 2b**), which are generally sensitive to schedule, and suggesting that the effects of outcome in **Figure 2e** are observable because individual differences in “switch” behavior is normalized in that analysis. Mutual information analysis revealed that there were also no group differences in the amount of information held from one trial to the next (**Supplemental Figure 2c**). Calculating the probability of exploration via the hidden Markov model (**Figure 2b**, left) we found no group differences in the amount of exploration throughout the restless bandit task (**Supplemental Figure 2d**). In order to assess the likelihood that mice are choosing the optimal choice across trials we employed p(Best) to analyze whether mice are choosing the highest probability reward. There were no group differences in p(Best) across restless bandit testing (**Supplemental Figure 2e**). There were no group differences in p(Best) across restless bandit testing (**Supplemental Figure 2e**). In order to assess the likelihood that mice are choosing the optimal choice across trials we employed p(Best) to analyze whether mice are choosing the highest probability reward.

We have previously found that the location of nosepokes on the touchscreen associated with an explore or exploit choice labeled by the hidden Markov model differs by decision making state, allowing us to investigate motor action differences within both states (Mueller et al. 2025). We replicate this effect, however, no genotype or sex differences were apparent in these measures. Distance between touches was significantly smaller in exploit than explore in both euclidean (**Supplemental Figure 3a,3b**, p < 0.001; ꞵ_state_ = 26.121) and mahalanobis measurements (**Supplemental Figure 3c,3d**; p < 0.001; ꞵ_state_ = 10.250). We also saw that exploit choices were farther from the center of the touchscreen, suggestive of reduced deliberation (**Supplemental Figure 3e,3f**, p < 0.001; ꞵ_state_ = -75.2678).

**Figure 3:**
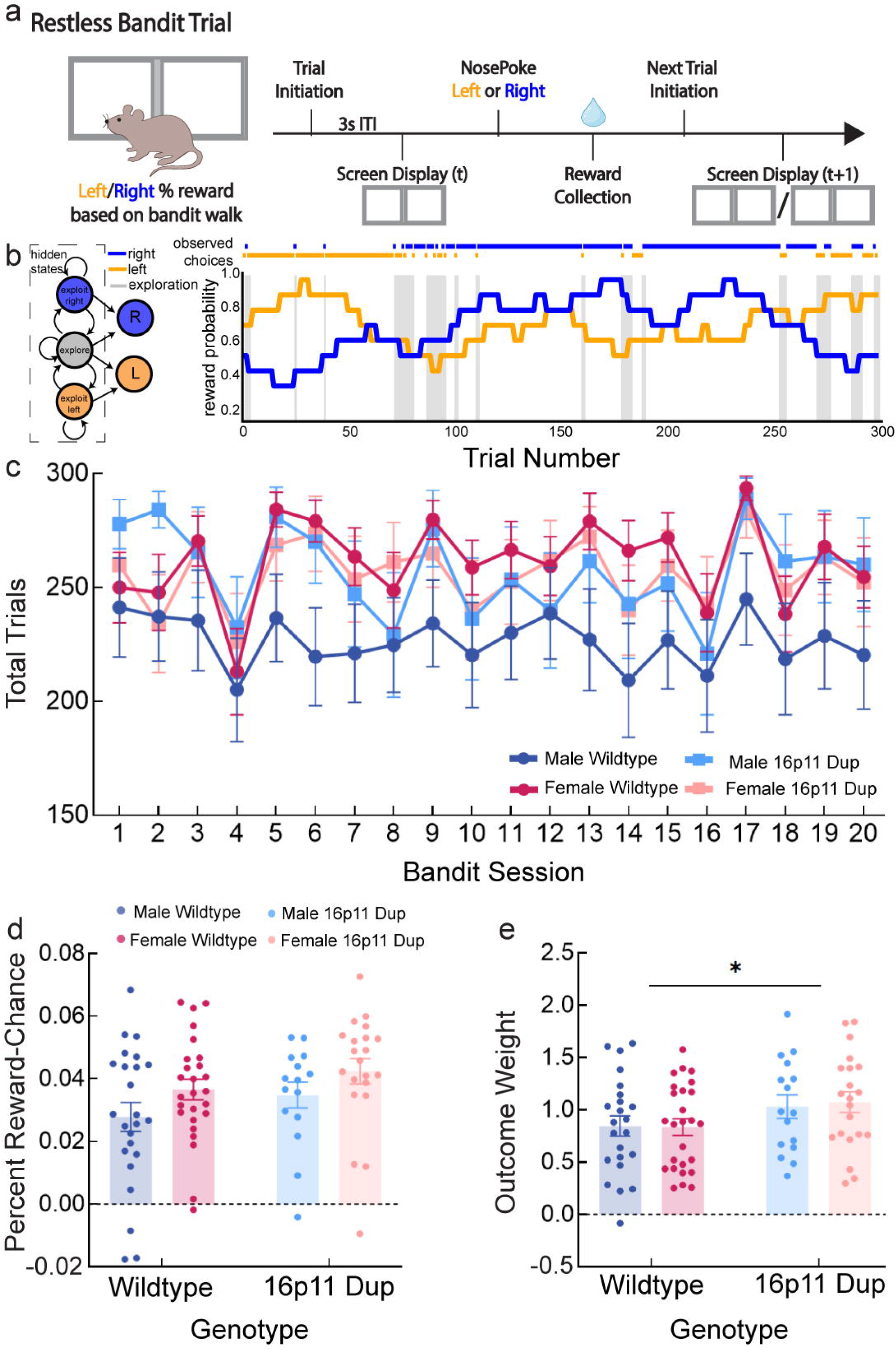
MK-801 prevents all animals from using past outcomes to select next choices. a) Schematic depicting the timeline of pharmacological manipulation in relation to bandit schedule, where alternating saline and MK-801 injections were given Tuesday through Friday for consecutive weeks. b) Averaged Percent Reward-Chance split by sex, genotype, and treatment. MK-801 significantly reduced Reward-Chance. c) Percent Reward-Chance displayed across restless bandit sessions 29-40 split by sex and genotype. Purple shading indicates MK-801 injection days, while no shading indicates saline injection days. MK-801 significantly reduced Reward-Chance across all groups. Red circles indicate female wildtype, blue circles indicate male wildtype, light blue squares indicate male 16p11 Dup, and orange squares indicate female 16p11 Dup mice. d) Averaged choice reaction time split by sex, genotype, and treatment. MK-801 significantly reduced choice reaction time. e) Choice reaction time in seconds displayed across restless bandit sessions 29-40 split by sex and genotype. Purple shading indicates MK-801 injection days, while no shading indicates saline injection days. MK-801 significantly reduced choice reaction time across all groups. Red circles indicate female wildtype, blue circles indicate male wildtype, light blue squares indicate male 16p11 Dup, and orange squares indicate female 16p11 Dup mice. f) Averaged reward retrieval reaction time split by sex, genotype, and treatment. MK-801 significantly reduced reward retrieval time. g) Reward retrieval reaction time in seconds displayed across restless bandit sessions 29-40 split by sex and genotype. Purple shading indicates MK-801 injection days, while no shading indicates saline injection days. MK-801 significantly reduced reward retrieval time across all groups. Red circles indicate female wildtype, blue circles indicate male wildtype, light blue squares indicate male 16p11 Dup, and orange squares indicate female 16p11 Dup mice. h) Percent Win-Stay split by genotype and sex. White indicates saline manipulation and purple indicates MK-801 manipulation. MK-801 significantly reduced Win-Stay behavior. i) Percent Lose-Shift split by genotype and sex. White indicates saline manipulation and purple indicates MK-801 manipulation. MK-801 had no effect on Lose-Shift behavior. j) Probability of exploration split by genotype and sex. White indicates saline manipulation and purple indicates MK-801 manipulation. MK-801 had no effect on the probability of exploration. k) Probability of choosing the highest probability choice split by genotype and sex. White indicates saline manipulation and purple indicates MK-801 manipulation. MK-801 significantly reduced the probability of choosing the correct choice. l) Outcome weight split by genotype and sex. White indicates saline manipulation and purple indicates MK-801 manipulation. MK-801 significantly reduced outcome weight. m) Mutual Information split by genotype and sex. White indicates saline manipulation and purple indicates MK-801 manipulation. MK-801 significantly reduced mutual information. For simplicity of visualization, all bar plots are averages across trials and sessions, so that each individual data point plotted represents the overall average for a mouse. Significant throughout this figure is represented in the following way: * p value less than 0.05. Error bars represent standard error of the mean (SEM).

### MK-801 prevents all animals from using past outcomes to select next choices

To examine the effect of NMDAR disruption on flexible decision making in the two-arm spatial restless bandit task, we used the NMDAR antagonist, MK-801 (Kruk-Slomka et al. 2016; Bygrave et al. 2016; Mandillo et al. 2003; Uehara et al. 2010; Stefani and Moghaddam 2005; Li et al. 2015). All animals received alternating days of saline and MK-801 systemic injections four days a week in a within-subjects design (**Figure 3a**). MK-801 impacted the ability to flexibly switch between choices to obtain the optimal amount of reward in all animals. MK-801 treatment significantly reduced percent reward-chance (**Figure 3b,3c**, p = 0.005; ꞵ_drug_ = 3.565) suggesting that under MK-801 treatment mice are earning significantly less reward as they are less successful at the two-arm restless bandit task. There is no overall sex or genotype difference on percent reward-chance.

MK-801 treatment also speeded reaction times. MK-801 treatment significantly reduced choice reaction time (**Figure 3d,3e**, p < 0.001; ꞵ_drug_ = -4.801) and reward retrieval reaction time (**Figure 3f,3g**, p = 0.005; ꞵ_drug_ = 3.565). There is no overall sex or genotype difference on either reaction time. Together these results suggest that MK-801 negatively impacts task accuracy in conjunction with speeding responses. Typically, reaction times decrease with *increased* confidence or certainty in decisions (Nissen and Bullemer 1987; Graybiel 2008), suggesting that NMDAR function under normal conditions supports the correlation between response vigor and reward success. This raises the question of how NMDAR antagonism with MK-801 influences animals’ choices so that they are less successful at getting rewards.

We hypothesized that MK-801 treatment would result in more shifts, similar to what is seen in humans experiencing psychosis and schizophrenia (Gold et al. 2008; Juckel et al. 2006; James A. Waltz et al. 2020; Schlagenhauf et al. 2014; Reed et al. 2020). MK-801 treatment significantly reduced percent Win-Stay compared to control (**Figure 3h**, p = 0.005; ꞵ_drug_ = 3.561) suggesting increased shifting behavior (switching choices) even following a rewarded choice. There is no overall sex or genotype difference on Win-Stay. MK-801 treatment had no effect on Lose-Shift (**Figure 3i**). Together this suggests that MK-801 might dull the influence of rewarded outcomes on subsequent trial choices, making mice shift when they should stay with a rewarded choice.

We next employed a set of modeling and computational approaches to understand what influences choices in animals under NMDAR antagonism. MK-801 treatment had no effect on the probability of exploration as assigned by the hidden Markov model (Chen, Knep, et al. 2021) suggesting that MK-801 is not affecting the likelihood of exploiting choices, but may reduce how optimal exploited choices are (**Figure 3j**). In line with this, MK-801 treatment significantly reduced the probability of animals making the “best” or optimal choice on any given trial (p(Best); **Figure 3k**, p = 0.014; ꞵ_drug_ = 2.975). Likewise, MK-801 treatment significantly reduced outcome weight on switching behavior (**Figure 3l**, p = 0.001; ꞵ_drug_ = 4.410) suggesting that outcomes are less influential on the decision to stay or switch on the next trial under the influence of MK-801. Similarly, MK-801 treatment significantly reduced mutual information (**Figure 3m**, p = 0.040; ꞵ_drug_ = 2.368), calculated as the ability to predict what the animal would choose on the next trial as a function of previous trial choice and outcome - in case there was a more complex relationship driving choice than a simple Win-Stay/Lose-Shift. Together, these results suggest that MK-801 prevents mice from using rewards to select optimal choices, but not from making repeated sequences of (less rewarding) choices.

## Discussion

This study tested decision making strategy among two models for risk factors for psychosis: 16p11.2 duplication (16p11 Dup) genetic mouse model and the NMDAR antagonist, MK-801. The first major finding is that early operant training reveals some male-biased impacts of 16p11 Dup, not dissimilar to those previously shown in mice modeling the inverse copy number variation, 16p11.2 hemideletion (N. M. Grissom et al. 2017). The second major finding is that MK-801 treatment reduced flexible switching between choices to obtain the optimal amount of reward during probabilistic decision making.

We saw male-biased impacts across a set of early operant training schedules, sometimes showing differential impacts on male wildtypes versus 16p11 Dup. However, the direction of the effect - whether wildtype males were the most engaged, or the least engaged group - differed across training. In particular, the introduction of two choices in the *100-0 deterministic* training reduced engagement of wildtype males that had previously been high, but did not impact 16p11 Dup males. This suggests that the strategies for learning, such as the degree of exploratory or off-target behavior engaged during uncertainty, may be a factor that is impacted by 16p11 Dup in males. The overall finding of some male specific impacts of 16p11 Dup is consistent with prior work showing male biased impacts of 16p11.2 hemideletion (N. M. Grissom et al. 2017; Angelakos et al. 2017; Kim et al. 2024; Lynch et al. 2020; Agarwalla et al. 2020; Kumar et al. 2018; Rojas, Heller, and Grissom 2023), and broadly consistent with other work showing male-biased impacts of neurodevelopmental disorder-linked genotypes (Chase et al. 2025). Many of these phenotypes have been identified in a learning and/or reward context, suggesting that sex differences in the function and/or development of neural circuits important for learning may be a critical factor in whether gene variants are associated with cognitive differences (N. Grissom et al. 2024; Nicola M. Grissom and Reyes 2018; Orsini and Setlow 2017). These mechanisms may potentially contribute to sex biases in diagnoses for neurodevelopmental and neuropsychiatric disorders (Lewine 1981; Lewine, Strauss, and Gift 1981; Aleman, Kahn, and Selten 2003).

Though 16p11 Dup is linked with psychosis which has been associated with altered decision making (Gold et al. 2008; Juckel et al. 2006; James A. Waltz et al. 2020; Schlagenhauf et al. 2014; Reed et al. 2020), we did not see overwhelming differences in restless bandit performance across genotypes in the current study, with the only effect being *increased* outcome sensitivity in 16p11 Dup. There are several possible reasons for this. First, the fact that we do see some 16p11 Dup effects in early operant training argues that novelty in the task, or early schema learning, may better expose behavioral phenotypes, whereas daily testing with a restless, but fundamentally similar task, does not engage the affected circuits. This is consistent with prior findings of impaired novelty-associated behaviors in 16p11.2 hemideletion (Yang et al. 2015). If so, tasks that explicitly require novel associations or rules, such as operant set shifting (Glewwe et al. 2025), may reveal cognitive impacts of 16p11 Dup. Additionally, 16p11.2 copy number variants and patients with schizophrenia have shown some evidence for aging effects or delayed onset cognitive effects (Qureshi et al. 2014; Harris and Jeste 1988; Howard et al. 2000), suggesting that future experimenters should implement behavioral testing across different age groups.

We next tested the NMDAR antagonist MK-801 to determine how it impacted decision making. NMDAR antagonists can cause psychosis-like experiences and cognitive deficits (Zick et al. 2018; MacDonald and Chafee 2006; Kummerfeld et al. 2020), and are thought to model reductions of NMDA-receptors found in the frontal cortex in schizophrenia (Vinogradov et al. 2023). MK-801 had large impacts on behavior, but did not exert these in a genotype specific manner. Future work may determine if this is a function of dose response, and that a lower or subthreshold dose of MK-801 may impact behavior in mice modeling psychosis relevant genotypes, while not impacting wildtypes. However, with the dose selected based on prior literature, we found that MK-801 manipulation introduces several deficits in two-arm spatial restless bandit task performance while decreasing overall response times. MK-801 affects the ability to flexibly switch between choices to obtain the optimal amount of reward. MK-801 treatment resulted in a lower tendency to stay with a rewarded choice, lower probability of selecting the highest rewarded option, and less influence of prior outcomes on future choices. This strongly parallels findings of altered decision making deficits in psychosis, including inconsistent choice selections (Geana et al. 2022; Pratt et al. 2021), increased “switching,” or exploration, between choices, and decreased selection of the highest value choice (Gold et al. 2008; Juckel et al. 2006; James A. Waltz et al. 2020; Schlagenhauf et al. 2014; Reed et al. 2020). Further study of MK-801 and other NMDAR antagonist impacts on cognition may therefore pay substantial dividends to understanding the pathophysiology behind altered cognition in psychosis.

Copy number variations affecting 16p11.2 have been associated with a host of neurodevelopmental/neuropsychiatric diagnoses, including but not limited to schizophrenia, autism, epilepsy/seizures, microcephaly, speech and language delay, depression, ADHD, and bipolar disorder (Rein and Yan 2020). The diversity of diagnoses suggests that while 16p11.2 copy number variation exists as a risk factor, it likely interacts with other factors to lead to a particular diagnosis, or any diagnosis at all. Understanding altered learning and decision making computations as a result of 16p11.2 copy number variants can help reveal crucial mechanisms driving vulnerability in clinical populations.

---

**Supplemental Figure 1.**
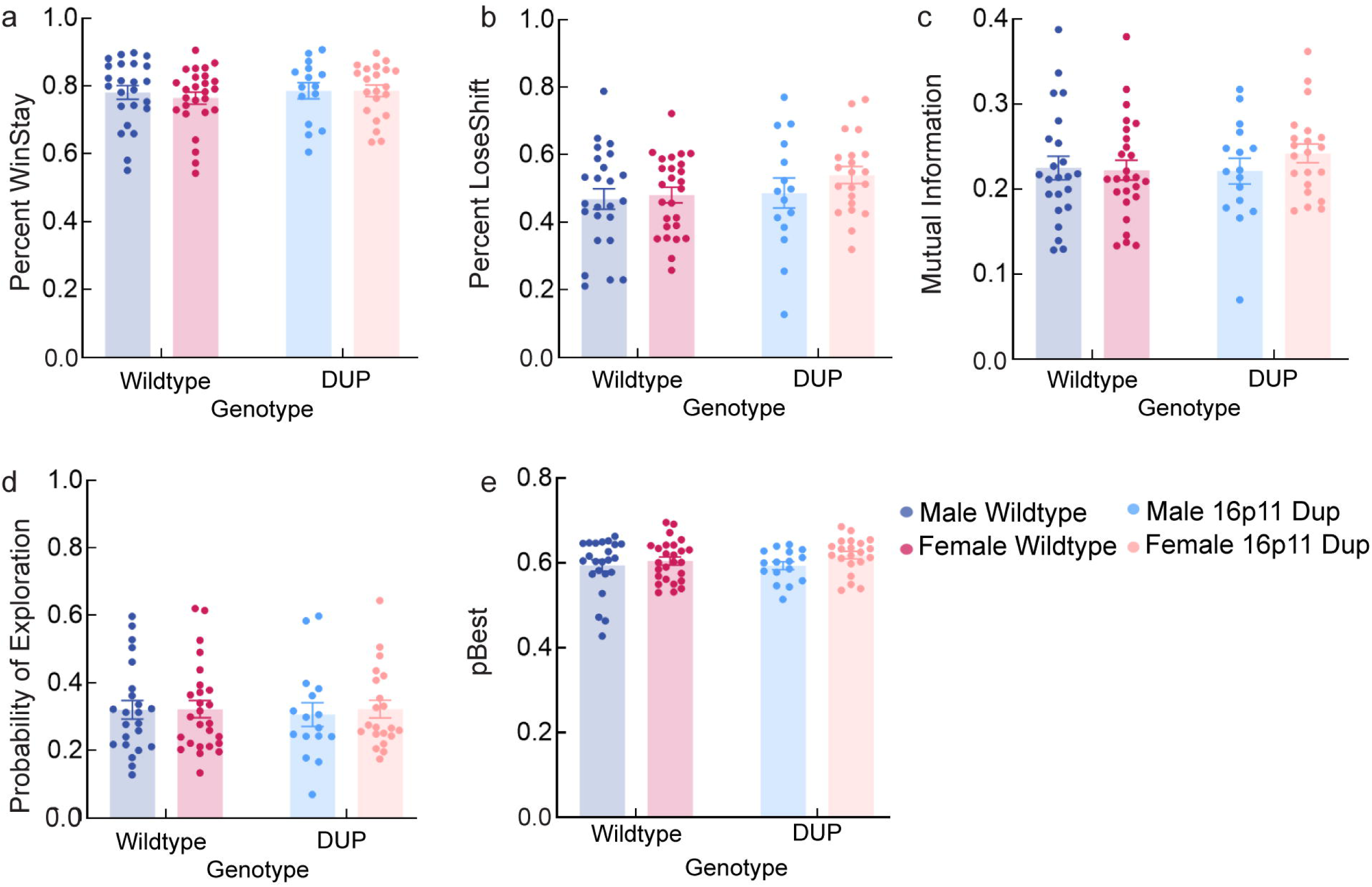
16p11.2 duplication prevents male-typical engagement with operant training, with no impact on females. a) Schematic illustrating chamber event calculations for choice reaction time, reward retrieval reaction time, and trial initiation reaction time measures. b) Schematic illustrating possible screen display and trial events for *Must Initiate* training schedule. c) Total trials displayed across the first 5 days of *Must Initiate* training split by sex and genotype. Males completed significantly more total trials. Planned pairwise comparisons revealed that male wildtypes complete more trials than all females on days 5 (all p ≤ 0.037; female wildtype indicated by “b”; female 16p11 Dup indicated by “c”). d) Choice reaction time displayed across the first 5 days of *Must Initiate* training split by sex and genotype. No significant differences between groups. e) Reward retrieval reaction time displayed across the first 5 days of *Must Initiate* training split by sex and genotype. No significant differences between groups. f) Trial Initiation reaction time displayed across the first 5 days of *Must Initiate* training split by sex and genotype. No significant differences between groups. g) Schematic illustrating possible screen display and trial events for *Punish Incorrect* training schedule. h) Total trials displayed across the first 5 days of *Punish Incorrect* training split by sex and genotype. No significant differences between groups. i) Choice reaction time displayed across the first 5 days of *Punish Incorrect* training split by sex and genotype. No significant differences between groups. j) Reward retrieval reaction time displayed across the first 5 days of *Punish Incorrect* training split by sex and genotype. No significant differences between groups. k) Trial Initiation reaction time displayed across the first 5 days of *Punish Incorrect* training split by sex and genotype. No significant differences between groups. For simplicity of visualization, all plots are averages across trials and mice for each session. Red circles indicate female wildtype, blue circles indicate male wildtype, light blue squares indicate male 16p11 Dup, and orange squares indicate female 16p11 Dup mice. Significance throughout this figure is represented in the following way: “a” indicates male wildtype significantly different from male 16p11 Dup, “b” indicates male wildtype significantly different from female wildtype, “c” indicates male wildtype significantly different from female 16p11 Dup. Error bars represent standard error of the mean (SEM).

**Supplemental Figure 2.**
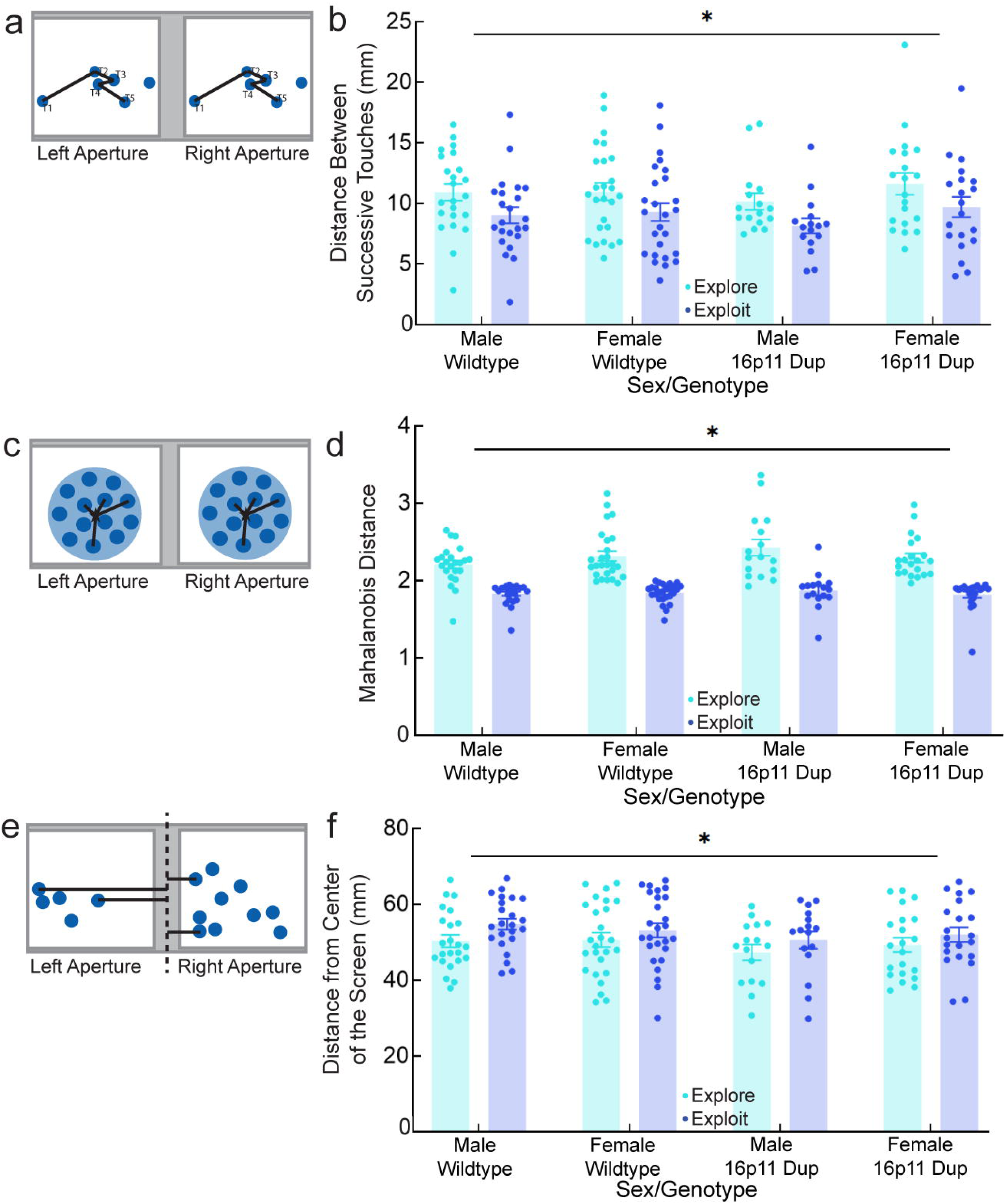
16p11 Duplication restless bandit performance extended. a) Percent Win-Stay split by genotype and sex. No significant differences between groups. b) Percent Lose-Shift split by genotype and sex. No significant differences between groups. c) Mutual Information split by genotype and sex. No significant differences between groups. d) Probability of Exploration as defined by HMM split by genotype and sex. No significant differences between groups. e) Probability of choosing the highest probability choice split by genotype and sex. No significant differences between groups. Red indicates female wildtype, blue indicates male wildtype, light blue indicates male 16p11 Dup, and orange indicates female 16p11 Dup mice. For simplicity of visualization, all plots are averages across trials and sessions, so that each individual data point plotted represents the overall average for a mouse. Significant throughout this figure is represented in the following way: * p value less than 0.05. Error bars represent standard error of the mean (SEM).

**Supplemental Figure 3:**
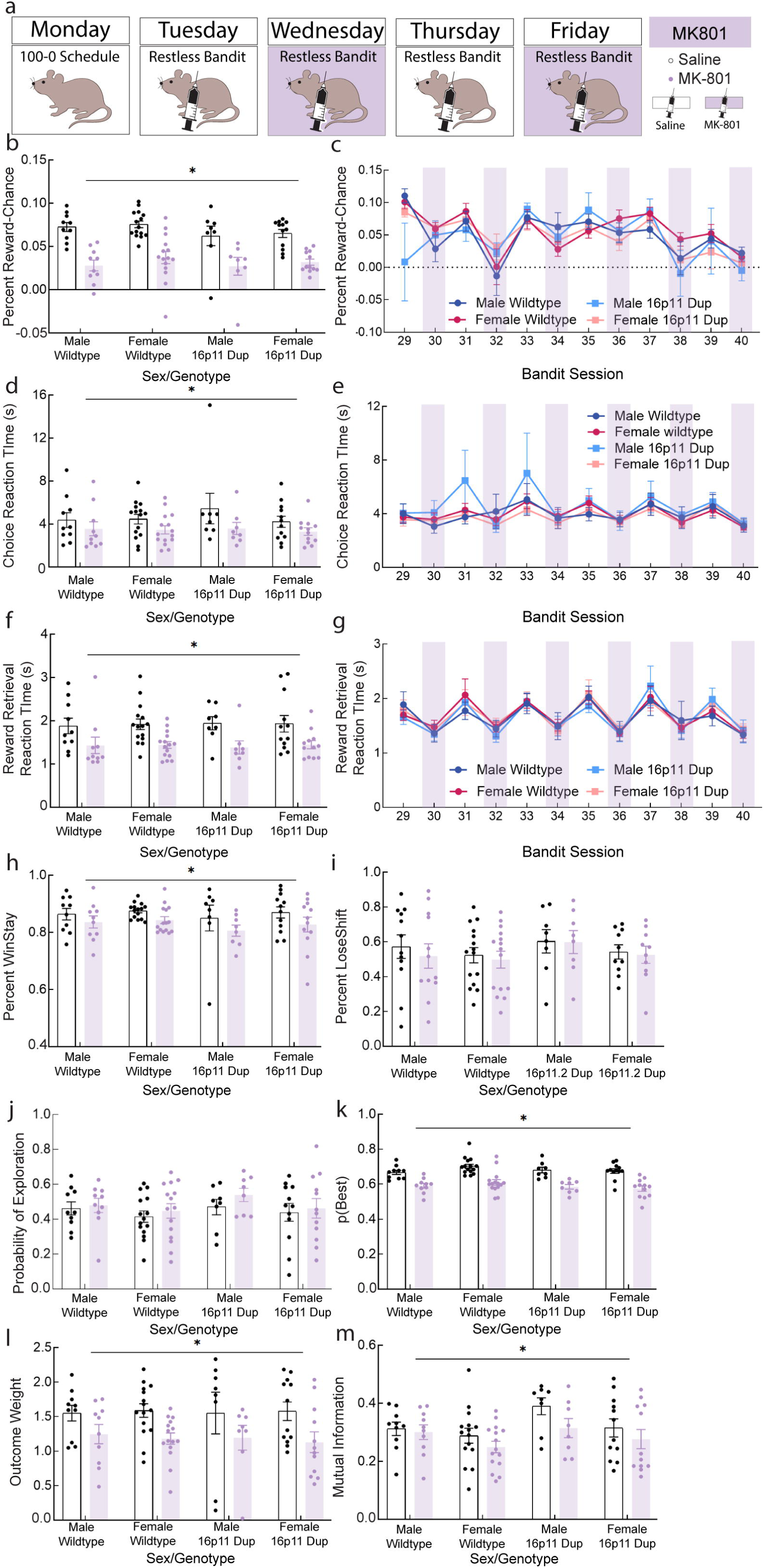
Exploit states reduce action variability during decision making regardless of sex and genotype. a) Schematic of Euclidean distance where the distance is calculated between touch 1 and touch 2, touch 2 and touch 3, touch 3 and touch 4, and so on. Shown here are possible left/right touches in blue and the distance relationship from one to another represented by black lines. b) Average Euclidean distance split by state, sex, and genotype. Exploit touches significantly reduced Euclidean distance. Light blue indicates distance between explore touches and dark blue indicates distance between exploit touches. c) Schematic of Mahalanobis distance where the individual data points are measured from the overall centroid of the dataset. Shown here are possible left/right Mahalanobis clusters (light blue circles) and centroids (stars) and the Mahalanobis distance relationship from each touch (darker blue circles) in a cluster to the centroid represented by black lines. d) Average Mahalanobis distance split by state, sex, and genotype. Exploit touches significantly reduced Mahalanobis distance. Light blue indicates Mahalanobis distance between explore touches and dark blue indicates Mahalanobis distance between exploit touches. e) Schematic of distance from the center of the screen where touch distance from both left and right choice apertures is measured from the midpoint of the operant screen. Shown here are possible left/right touches in blue and the distance of each from the center of the touchscreen represented by black lines. f) Average distance from the center of the screen split by state, sex, and genotype. Explore touches were significantly closer to the center of the screen. Light blue indicates distance from the center of the screen for explore touches and dark blue indicates distance from the center of the screen for exploit touches. For simplicity of visualization, all plots are averages across trials and sessions, so that each individual data point plotted represents the overall average for a mouse. Significant throughout this figure is represented in the following way: * p value less than 0.05. Error bars represent standard error of the mean (SEM).

## Acknowledgments

This work was supported by NIMH R01 MH123661 (NMG), NIMH P50 MH119569 (NMG, PER, SRH), Canada Research Chair Dynamics of Cognition FD507106 (RBE), and NIMH T32 training grant MH115886 (DM). Thank you to Nic Glewwe and Micaela Porod for helping improve this manuscript. Thank you to lab staff A Yang, Kira Stetler, and Laura Garbe, along with all our wonderful undergraduate students for running daily behavioral tasks.

